## Supplementary files and figures for "High resolution analysis of proteome dynamics during *Bacillus subtilis* sporulation": Supplementary figures.pdf

### High resolution time-lapse proteomic analysis of *Bacillus subtilis* sporulation

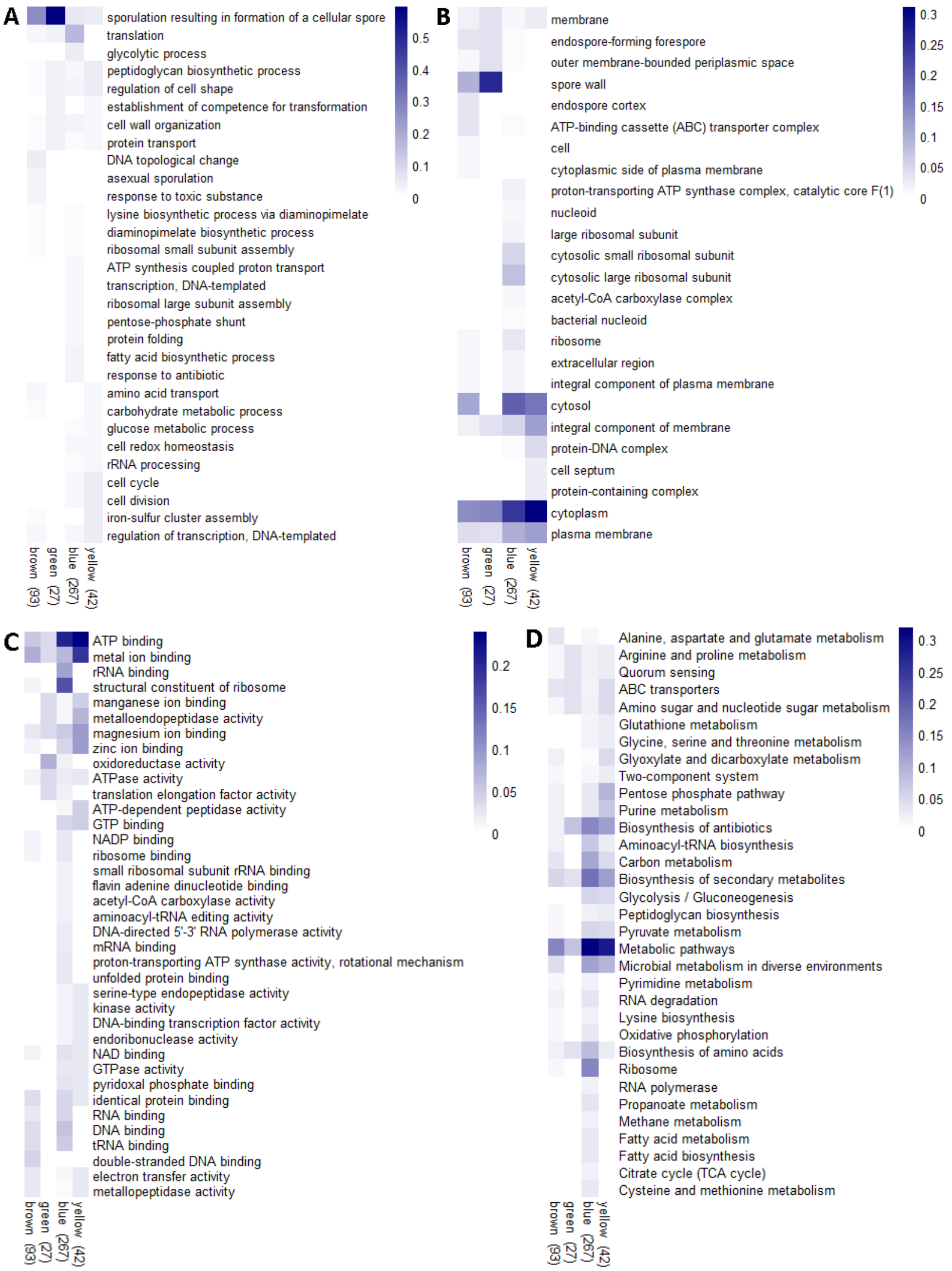

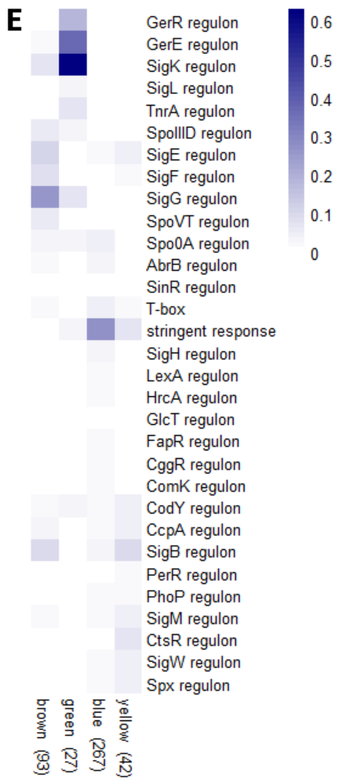


**Figure S1. Heatmap visualization of GO (gene ontology) annotation (A, B and C), KEGG pathway (D) and transcriptional regulons (E).** (A) GO biological process. (B) GO cellular compartments. (C) GO molecular functions. The progressively darker blue indicates the fractions of the module proteins belongs to the categories labeled on the right of the heatmap. GO categories, KEGG pathways and Regulons with more than 3 proteins in total were shown in the heatmap. Module size is displayed in the brackets flowing the module name.
