## Supplementary files and figures for "High resolution analysis of proteome dynamics during *Bacillus subtilis* sporulation": data imputation.R

```
#########imputation with raw data###########################
#install.packages("Amelia")
library(Amelia)
SetDir <- choose.dir() #run this and you will get a popup window to choose the directory where your data is.
setwd(SetDir) # this then sets the working directory to your data directory.
data <- read.table("time series with three data.txt", header = TRUE, sep = "\t", na.strings = "A")
rownames(data) <- data$A
data1 <- data[,2:70]
data1 <- t(data1)

n_value <- data[,71]
data2 <- data1[,n_value >=46]
length(which(n_value ==69))
length(which(n_value >=46))
#timepoint
timepoint <- as.numeric(c(
  rep(seq(from = 0, to = 180, by = 15),rep(3,13)),
  rep(seq(from =210, to = 480, by = 30),rep(3,10))
)
)
#replicate
replica <- rep(1:3,23)
#log transformation
data2 <- log2(data2)

#imputation

a <- cbind(data2, timepoint, replica)

a.out.time <- amelia(a, m =1, ts = "timepoint", cs = "replica", polytime = 2,intercs = TRUE, empri = 10000, p2s = 0)
missmap(a.out.time)

write.amelia(obj=a.out.time, file.stem = "time series with three log2 data_missing data imputed_polytime_two") #m = 1


#missingness map*********************************************************************************
data_2 <- data1
data_2[is.na(data_2)==F] <- 1
data_2[is.na(data_2)==T] <- 0
t <- c(
  seq(from = 1, to = 67, by = 3),
  seq(from = 2, to = 68, by = 3),
  seq(from = 3, to = 69, by = 3)
)
data_2 <- data_2[t,]

library(pheatmap)

rownames(annotation_col) = colnames(data_2)

annotation_row = data.frame(
  Replication = factor(rep(c("Replicate 1", "Replicate 2", "Replicate 3"), c(23, 23, 23)))
)
rownames(annotation_row) = rownames(data_2)
pheatmap(data_2,color = colorRampPalette(c("gray", "darkgreen"))(2), cluster_rows = F, cluster_cols = F, show_rownames=F, legend_breaks = 0:1, legend_labels = c("Missing","Observed"),
         show_colnames = F, annotation_row = annotation_row, legend = F, 
          border_color=NA, fontsize = 10, 
         fontsize_row = 10, height=20)
```
