## Supplementary files and figures for "High resolution analysis of proteome dynamics during *Bacillus subtilis* sporulation": rcode-WGCNA_.R

```
#BiocManager::install("WGCNA")
SetDir <- choose.dir() #run this and you will get a popup window to choose the directory where your data is.
setwd(SetDir) # this then sets the working directory to your data directory.
# Load the WGCNA package
library(WGCNA);
# The following setting is important, do not omit.
options(stringsAsFactors = FALSE);
#Read in the female liver data set
femData1 = read.csv("time series with three log2 data_missing data imputed.csv", header = T)
 
rownames(femData1) <- femData1[,1]

femData <- femData1[,-c(1,431,432)]

# Take a quick look at what is in the data set:
dim(femData)
names(femData)

#average data
data2 <- femData[1:2,]
m <- 1
while(m<=23){
        i <- 3*m
        j <- i-2
        data2[m,] <- colMeans(femData[j:i,])
        m <- m+1
}
rownames(data2) <-  as.numeric(c(
        seq(from = 0, to = 180, by = 15),
        seq(from =210, to = 480, by = 30)
)
)
t <-  as.numeric(c(
        seq(from = 0, to = 180, by = 15),
        seq(from =210, to = 480, by = 30)
)
)
 
datExpr = as.data.frame(data2)
nGenes = ncol(datExpr)
nSamples = nrow(datExpr)

# Choose a set of soft-thresholding powers
powers = c(c(1:10), seq(from = 12, to=40, by=2))
# Call the network topology analysis function
sft = pickSoftThreshold(datExpr, powerVector = powers, verbose = 5, networkType = "signed")
# Plot the results:
sizeGrWindow(9, 5)
par(mfrow = c(1,2));
cex1 = 0.9;
# Scale-free topology fit index as a function of the soft-thresholding power
plot(sft$fitIndices[,1], -sign(sft$fitIndices[,3])*sft$fitIndices[,2],
     xlab="Soft Threshold (power)",ylab="Scale Free Topology Model Fit,signed R^2",type="n",
     main = paste("Scale independence"));
text(sft$fitIndices[,1], -sign(sft$fitIndices[,3])*sft$fitIndices[,2],
     labels=powers,cex=cex1,col="red");
# this line corresponds to using an R^2 cut-off of h
abline(h=0.80,col="red")
# Mean connectivity as a function of the soft-thresholding power
plot(sft$fitIndices[,1], sft$fitIndices[,5],
     xlab="Soft Threshold (power)",ylab="Mean Connectivity", type="n",
     main = paste("Mean connectivity"))
text(sft$fitIndices[,1], sft$fitIndices[,5], labels=powers, cex=cex1,col="red")
###
softPower = 9;
adjacency = adjacency(datExpr, type = "signed",power = softPower);

# Turn adjacency into topological overlap
TOM = TOMsimilarity(adjacency);
dissTOM = 1-TOM
# Call the hierarchical clustering function
geneTree = hclust(as.dist(dissTOM), method = "average");
# Plot the resulting clustering tree (dendrogram)
sizeGrWindow(12,9)
plot(geneTree, xlab="", sub="", main = "Gene clustering on TOM-based dissimilarity",
     labels = FALSE, hang = 0.04)
# We like large modules, so we set the minimum module size relatively high:
minModuleSize = 15;
# Module identification using dynamic tree cut:
dynamicMods = cutreeDynamic(dendro = geneTree, distM = dissTOM,
                            deepSplit = 2, pamRespectsDendro = FALSE,
                            minClusterSize = minModuleSize);
table(dynamicMods)
# Convert numeric lables into colors
dynamicColors = labels2colors(dynamicMods)
table(dynamicColors)
# Plot the dendrogram and colors underneath
sizeGrWindow(8,6)
plotDendroAndColors(geneTree, dynamicColors, "Dynamic Tree Cut",
                    dendroLabels = FALSE, hang = 0.03,
                    addGuide = TRUE, guideHang = 0.05,
                    main = "Gene dendrogram and module colors")
# Calculate eigengenes
MEList = moduleEigengenes(datExpr, colors = dynamicColors)
MEs = MEList$eigengenes
# Calculate dissimilarity of module eigengenes
MEDiss = 1-cor(MEs);
# Cluster module eigengenes
METree = hclust(as.dist(MEDiss), method = "average");
# Plot the result
sizeGrWindow(7, 6)
plot(METree, main = "Clustering of module eigengenes",
     xlab = "", sub = "")
MEDissThres = 0.25
# Plot the cut line into the dendrogram
abline(h=MEDissThres, col = "red")
# Call an automatic merging function
merge = mergeCloseModules(datExpr, dynamicColors, cutHeight = MEDissThres, verbose = 3)
# The merged module colors
mergedColors = merge$colors;
# Eigengenes of the new merged modules:
mergedMEs = merge$newMEs
plotMEpairs(mergedMEs)
sizeGrWindow(12, 9)
#pdf(file = "Plots/geneDendro-3.pdf", wi = 9, he = 6)
plotDendroAndColors(geneTree, cbind(dynamicColors, mergedColors),
                    c("Dynamic Tree Cut", "Merged dynamic"),
                    dendroLabels = FALSE, hang = 0.03,
                    addGuide = TRUE, guideHang = 0.05)
# Rename to moduleColors
moduleColors = mergedColors
# Construct numerical labels corresponding to the colors
colorOrder = c("grey", standardColors(50));
moduleLabels = match(moduleColors, colorOrder)-1;
MEs = mergedMEs;
#check clusters in a excel file if needed
datExpr1 <- rbind(datExpr, moduleColors, moduleLabels)
write.csv(datExpr1,"TOM cluster_softthreshold_signed.csv",)

# Transform dissTOM with a power to make moderately strong connections more visible in the heatmap
plotTOM = dissTOM^7;
# Set diagonal to NA for a nicer plot
diag(plotTOM) = NA;
plotTOM1 <- 1-plotTOM
# Call the plot function
sizeGrWindow(9,9)
TOMplot(plotTOM1, geneTree, moduleColors, main = "Network heatmap plot, all proteins")
#Export the network for Cytoscape

modules2 = c("brown", "green", "blue", "yellow")
# Select module probes
probes = names(datExpr)
inModule = is.finite(match(moduleColors, modules2));
modProbes = probes[inModule];

# Select the corresponding Topological Overlap
modTOM = TOM[inModule, inModule];

dimnames(modTOM) = list(modProbes, modProbes)
# Export the network into edge and node list files Cytoscape can read
cyt = exportNetworkToCytoscape(modTOM,
                               edgeFile = paste("CytoscapeInput-edges-threshold-0.2-cell-removed-soft-signed ", paste(modules2, collapse="-"), ".txt", sep=""),
                               weighted = TRUE,
                               threshold = 0.2,
                               nodeNames = modProbes,
                                                             nodeAttr = moduleColors[inModule])
####heatmap visualization####
library(pheatmap)

orderr <- order(moduleColors)
data2 <- datExpr[,orderr]

Z_data2 <- t(scale(data2))

#annotation row
annotation_row = data.frame(
        Cluster = factor(sort(moduleColors), levels = c("blue", "brown", "green", "yellow"))
)
row.names(annotation_row) = rownames(Z_data2)
#annotation colors
annotation_row$Cluster <- factor(annotation_row$Cluster, levels = c("blue", "brown", "green", "yellow"))
clustercol <- c("blue", "brown", "green", "yellow")
names(clustercol) <- levels(annotation_row$Cluster)
ann_colors = list(
        Cluster = clustercol
)
pheatmap(Z_data2,cluster_rows = F, cluster_cols = F, show_rownames=F, clustering_distance_rows="euclidean",
         clustering_distance_cols="euclidean", clustering_method="complete", show_colnames = T,
         annotation_row = annotation_row, annotation_colors = ann_colors, border_color=NA, fontsize = 10, 
         fontsize_row = 10, height=20)
```
